# *Rotiferometer*: an automated system for quantification of rotifer cultures

**DOI:** 10.1101/2025.05.12.653399

**Authors:** Alioune Diouf, Leandre Bereziat, David Nodier, Marco Amaral, Sinan Haliyo, Abdelkrim Mannioui

## Abstract

Accurate quantification and continuous monitoring of *Brachionus plicatilis L*. rotifer cultures are essential for aquaculture and aquatic animal research laboratories. Manual counting methods are labor-intensive, error-prone, and inefficient for large-scale operations, necessitating automated solutions. This study presents the Rotiferometer, an automated and cost-effective system that integrates mechanical design, deep learning, and automation for precise rotifer detection, classification and counting. Using a YOLOv8 model, the system achieves a mean average precision (mAP@0.5) of 94.7% in distinguishing gravid and non-gravid rotifers. It proceeds by scanning a 1 mL Sedgewick Rafter slide under 3 minutes, ensuring rapid and accurate enumeration. A strong correlation was observed between manual and Rotiferometer counts, (with R2 values of 0.9729 and 0.9868 for gravid and non-gravid rotifers, respectively), confirming the system s accuracy. Additionally, the analysis of operator variability using the Rotiferometer delivered consistent results regardless of the user, minimizing the need for specialized expertise. Finally, a 45-day monitoring experiment with the Rotiferometer effectively tracked rotifer population changes, identifying key phases of growth, decline, and recovery. These results highlight the device s potential to enhance rotifer culture management by providing real-time, reliable, and automated monitoring, thereby optimizing aquaculture productivity and research efficiency.

## INTRODUCTION

Rotifers, *Brachionus plicatilis L*., a microscopic zooplankton species approximately 180 μm in size, play a fundamental role in aquaculture and aquatic animal research laboratories (1). They serve as the primary live feed for fish larvae due to their high nutritional value, which includes essential proteins, lipids, and vitamins critical for larval development (2). This species is the most cultured due to its adaptability to controlled environments and rapid reproductive cycle (3). These characteristics make this species indispensable in hatcheries, especially in the larviculture of seabass (*Dicentrarchus labrax*), turbot (*Scophthalmus maximus*), and zebrafish (*Danio rerio*), where consistent and reliable live food availability is critical for survival and growth (4,5).

The semi-continuous culture method is widely used in aquaculture facilities to maintain stable rotifer populations. This technique involves daily harvesting, from 25% to 50%, and replenishment of the culture to balance population density and nutrient levels.

The indicator of active rotifers reproduction in the culture is the gravid rotifer, which are females carrying developed eggs inside their body. The gravid rotifers typically range from 25% to 50% of the population, with 25% being a sufficient ratio to double the population daily (6) Under optimal conditions, rotifer cultures can reach densities of up to 2.106 individuals per milliliter, with population doubling times typically exceeding 24 hours (1). However, maintaining culture stability poses significant challenges, including pathogen contamination, fluctuations in water quality, and accumulation of metabolic waste, which can lead to sudden population crashes (3).

Daily measurement of the number of rotifers in culture is therefore essential to ensure adequate monitoring of optimal culture (7). Traditional rotifer quantification methods rely on manual counting under a microscope, a labor-intensive and time-consuming process prone to human error. The process takes approximately 20 minutes per session and is often inaccurate, with results varying between operators.

A number of seminal studies have contributed to the development of automated rotifer counting systems. One study used a shape descriptor and an artificial neural network (ANN) to classify rotifers in iodine-fixed specimens at high magnification (x40). Another study introduced a computer vision-based system using frame-differencing background subtraction (FDBGS) for rotifer density detection and estimation. The system was then integrated into an automated fish larvae feeder that adjusted feeding rates according to rotifer density. In addition, Stelzer et al. (2009) used FDBGS to develop a rotifer sampling and counting system (8). They pursued by developing a prototype control system for rotifer cultures designed to monitor density and egg rates (9). The effectiveness of this system was validated using simulated data.

Recently, several methods have been developed for the detection and counting of rotifers, with a particular focus on Deep Learning (DL) techniques. A study using DL for automated tracking and counting of plankton, specifically YOLOv5 (You Only Look Once, version 5), including rotifers, demonstrated high accuracies with a higher Mean Average Precision (mAP) of 0.992% (10). A YOLOv3 model for rotifer detection in aquaculture environments was achieved with a mAP of approximately 93%, highlighting its effectiveness in high-density environments for counting rotifers to estimate the amount of rotifers for feeding or the density population in a rotifer culture (11). The model was tested with an average accuracy of 85.1% and an average inference speed of 1.4645 seconds. A recent study developed an improved YOLO model by incorporating deformable convolutional networks and an attention mechanism to enhance its detection capabilities (12). The improved YOLO model achieved a mAP of 93.7%, an improvement of 4.3% over the basic YOLOv5 model.

While progress has been made in the use of DL techniques for rotifer detection, the distinction between gravid (egg-bearing) and non-gravid (non-egg-bearing) rotifers remains unsolved. Indeed, distinguishing between gravid and non-gravid rotifers using deep learning (DL) techniques is a promising but yet underdeveloped. While advances in DL have facilitated species detection and classification, fine-grained tasks such as identifying physiological states (e.g., gravid vs. non-gravid) present opportunities.

Furthermore, counting strategies based on DL are proposed, but there is a paucity of literature on automated counting tools employing DL trained models to streamline the entire counting process, giving the tendencies, adjusting the culture or feeding. Alternatively, existing counting systems are prohibitively expensive due to the technologies employed.

To address these challenges, we have developed a new low-cost automated tool that uses X-Y axis scanning and image capture to cover the entire surface of a slide (Sedgewick Rafter) containing 1 ml of rotifers. In addition to accurately identifying gravid and non-gravid rotifers, this system allows for systematic counting of rotifers using a mechanical device, thereby facilitating automation and a trained DL model to expedite the counting process. This tool not only assesses the number of rotifers in the culture, but also stores data and generates real-time graphs showing the state and density of the culture. This allows precise adjustments to be made to the algae distribution and ensures accurate rotifer delivery to the larvae. The tool reduces counting errors, eliminates operator variability and frees the operator to focus on other tasks, particularly those related to animal welfare.

## Materials and Methods

### The Rotiferometer Design

The Rotiferometer is equipped with a motorized stage, a fixed digital microscope, lighting system, and image analysis software, has dimensions of 120 mm (width) × 175 mm (depth) × 200 mm (height), and weighs 0,645 kg. As illustrated in Figure 1.a, the servomotors S1 controls X-axis motion of the motorized stage via the larger gear, while the servomotor S2 controls Y-axis motion via the smaller gear. The carriage (See Figure 1.b) is adapted for a Sedgewick Rafter chamber 76 x 40 mm, which can move up to 5 cm along the X-axis and 2 cm along the Y-axis within the motion limit of the servo motors. Although the Rotiferometer is designed to scan a Sedgewick Rafter chamber, it can be easily adapted to other disposables.

**Figure 1.**
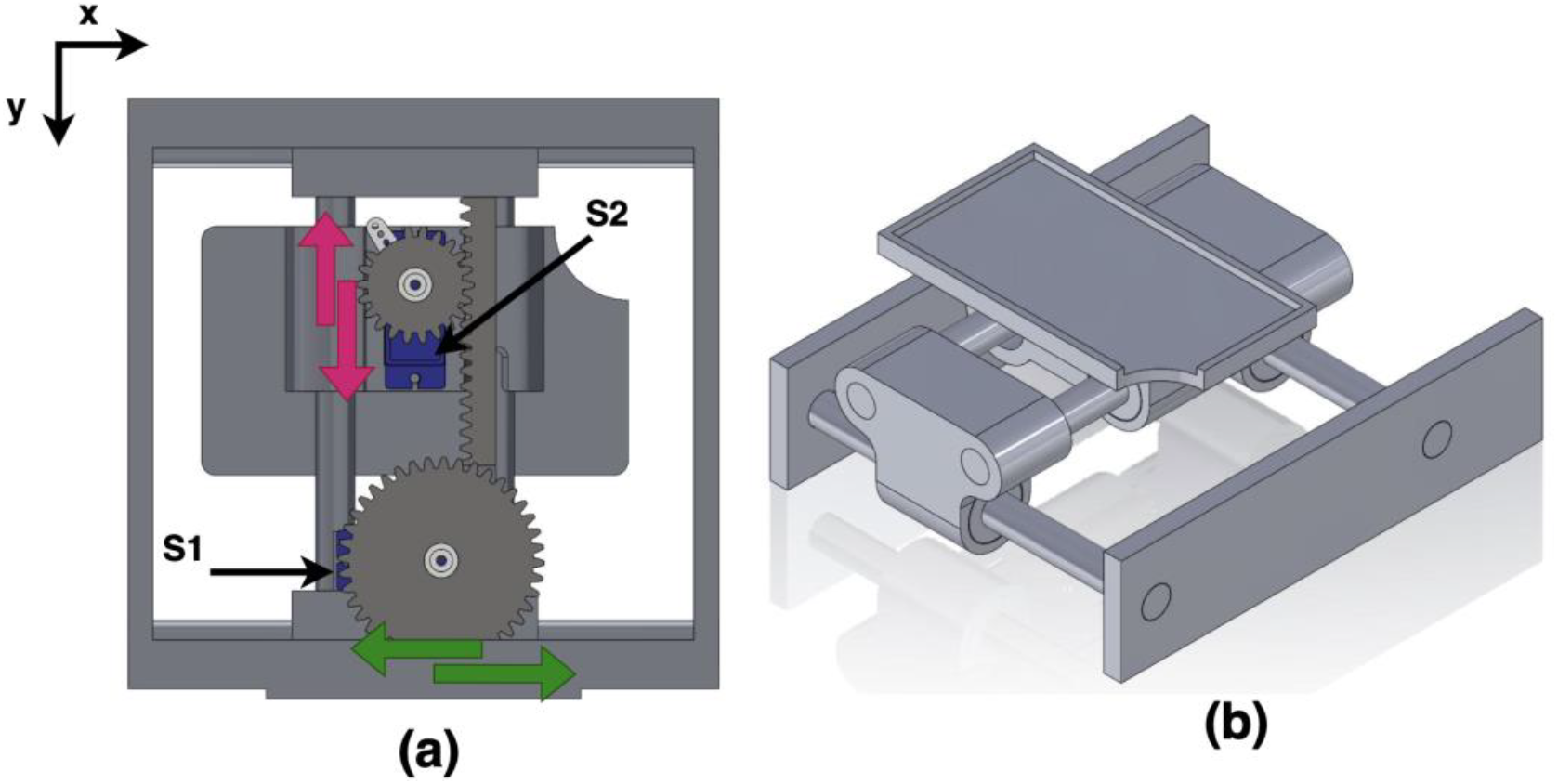
hardware design of the Rotiferometer. **(a)**: Bottom view, the rack and pinion system. **(b)**: Top view, prototype of carriage design.

Figure 2 shows an overview of the Rotiferometer system, showing its main hardware components, including the digital microscope, robotic arm, cables, plate and base. The microscope is a digital model designed for high-resolution imaging, allowing detailed visualization of specimens. The Swivelarm holding the microscope provides 1 degree of freedom, allowing positioning and focus adjustment. The cabling includes power cables for electronic components and data transfer. An ATmega 328p microcontroller was used to control servomotors and light. The plate provides a stable platform for holding specimens, with optional features such as adjustability and integrated lighting for improved visibility. The base provides robust support to minimize vibration and maintain stability during operation.

**Figure 2.**
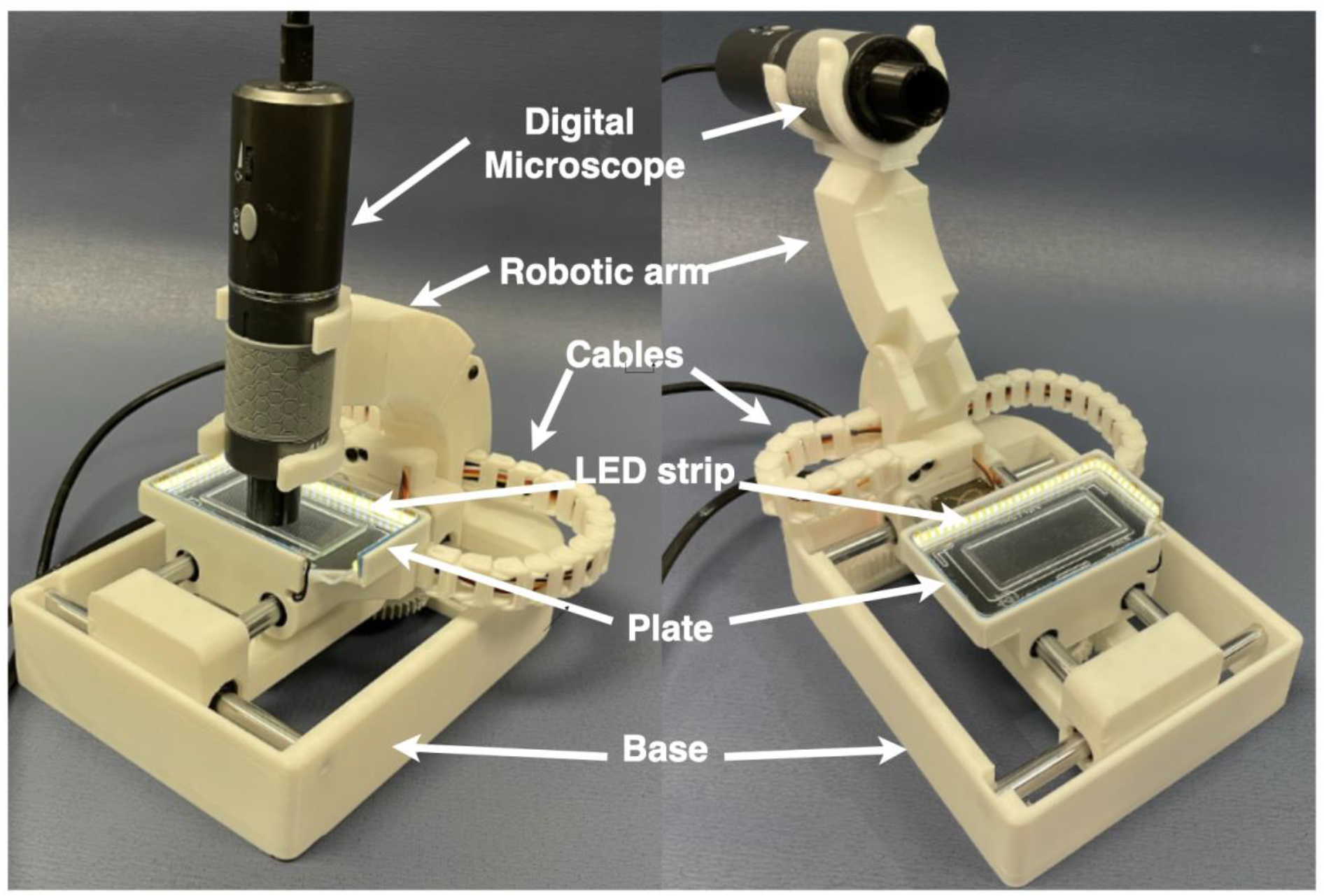
Overview of the Hardware design of the Rotiferometer.

### Lighting System

To ensure accurate rotifer detection, we replaced the built-in lighting system of the digital microscope with transmitted illumination, by installing an LED strip around the Rotiferometer stage (see Figure 2). Through experimentation with various light intensities, we found that low-intensity, in-line LED illumination provided high contrast, making rotifers appear bright against a dark background (Figure 3.a). This approach significantly reduced the visibility of contaminants, as shown in Figure 3.b. The LED strip is connected to a microcontroller, enabling precise intensity adjustments through software.

**Figure 3.**
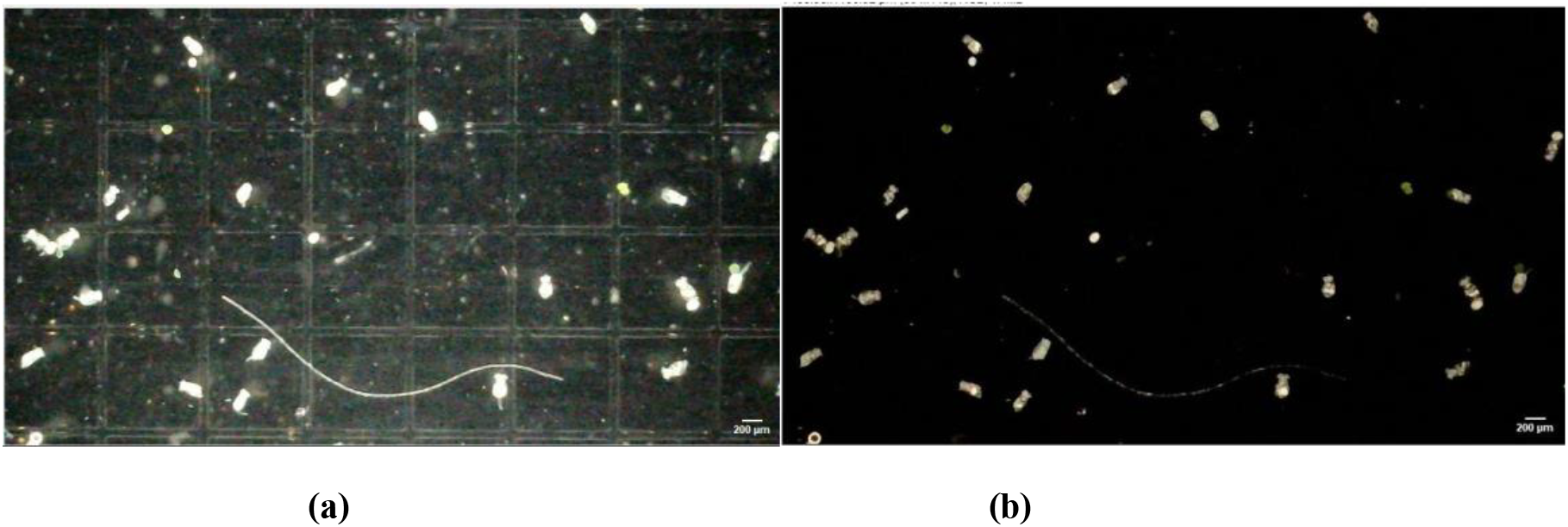
**(a) :** Coaxial lighting. **(b) :** Images captured with high and low intensity of transmitted lighting

### Rotifer culture

The rotifer, Brachionus plicatilis, was cultured in water with a salinity of 15 ppt, prepared by dissolving 15 g of Instant Ocean® sea salts in osmotic water under vigorous stirring. The rotifers were fed automatically using a timer-controlled peristaltic pump, which dispensing RotiGrow Nanno® algal concentrate stored at 4°C. The timer was programmed to deliver feed once per hour, and the system was checked daily to ensure a sufficient supply of algal concentrate and to prevent clogging in the supply lines. The rotifer culture was maintained at a room temperature of 25–26°C with continuous aeration. To maintain optimal culture conditions, the culture containers were replaced every two days.

### Rotifer Preparation

Rotifers from 1 ml of the culture were evenly distributed on a Sedgewick Rafter counting chamber, which measures 50 mm x 20 mm x 1 mm. To immobilize them during the process, they were treated with vinegar, which effectively stopped their movement.

### The Software

The software controls a digital microscope (with magnification ranging from 50 to 1000x), the two servo motors and the light intensity. The digital microscope captures a series of high-resolution images covering the standardized slide. Once the scanning is completed, the images are then processed by the deep learning algorithm trained on an accredited database to detect, identify, and enumerate the Rotifer (gravid and non-gravid). The software then calculates the concentration of different rotifer species in the total volume. The output data is processed to instantly display the results in graphical form and saved in CSV format (See Figure 4).

**Figure 4.**
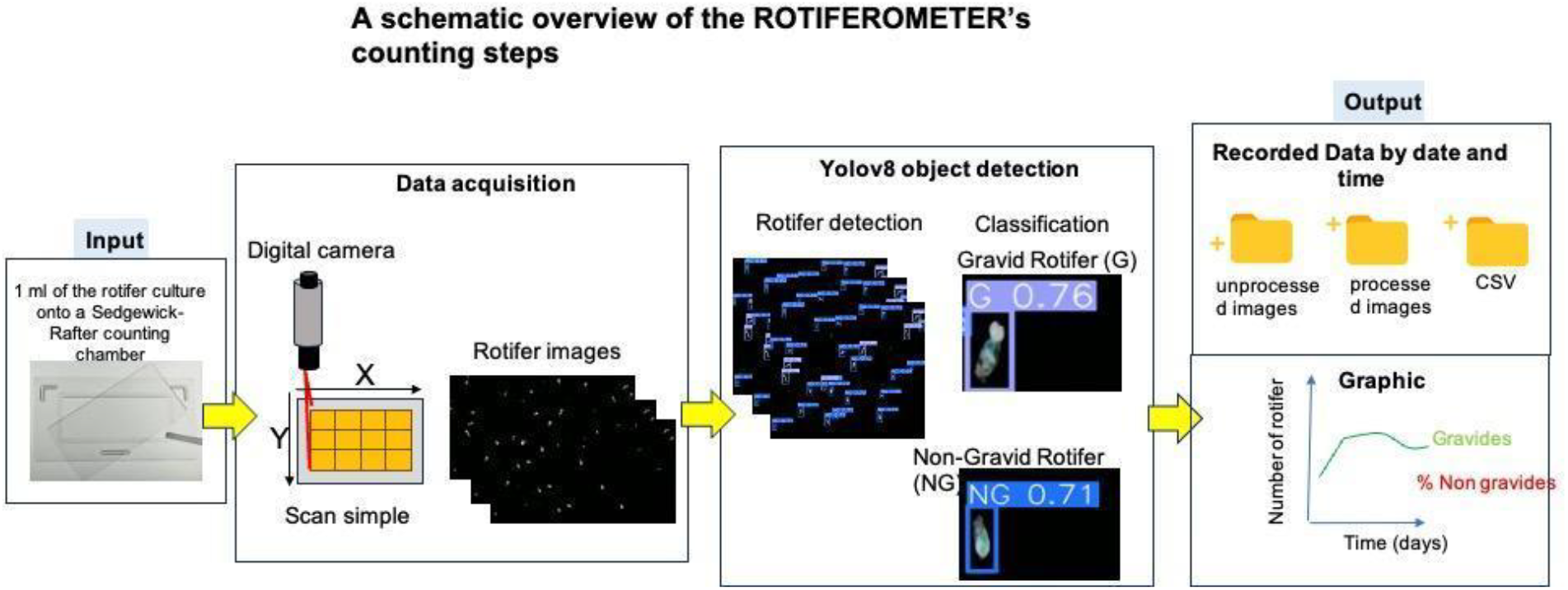
A streamlined method for estimating rotifer populations with the Rotiferometer

### Dataset collection, labelling

The DL algorithm is trained on a custom-made database, constructed from the images obtained from the same capture pipeline. A total of 642 images, comprising 7316 annotations, were utilized for analysis, as detailed in Table 1. The dataset was prepared using the online platform RoboFlow. Rotifers in each image were labeled into one of two categories: ‘*G*’ for gravid and ‘*NG*’ for non-gravid rotifers. To enhance the training, we focused on samples with low rotifer concentrations, resulting in images predominantly containing a small number of annotations (1–4 annotations per image; 831 images), as illustrated in Figure 5. Notably, the class distribution of ‘G’ and ‘NG’ reveals a significant imbalance, with the ‘*NG*’ class comprising over three times the number of samples compared to the ‘*G*’ class. This disparity reflects the naturally low prevalence of gravid individuals in the rotifer culture, as summarized in Table 1.

**Table 1.** Class balance on dataset.

| Classes | Train | Validation | Test | Total |
| --- | --- | --- | --- | --- |
| NG | 4118 | 1053 | 614 | 5785 |
| G | 1053 | 320 | 158 | 1531 |
| all | 5171 | 1373 | 772 | 7316 |

**Figure 5.**
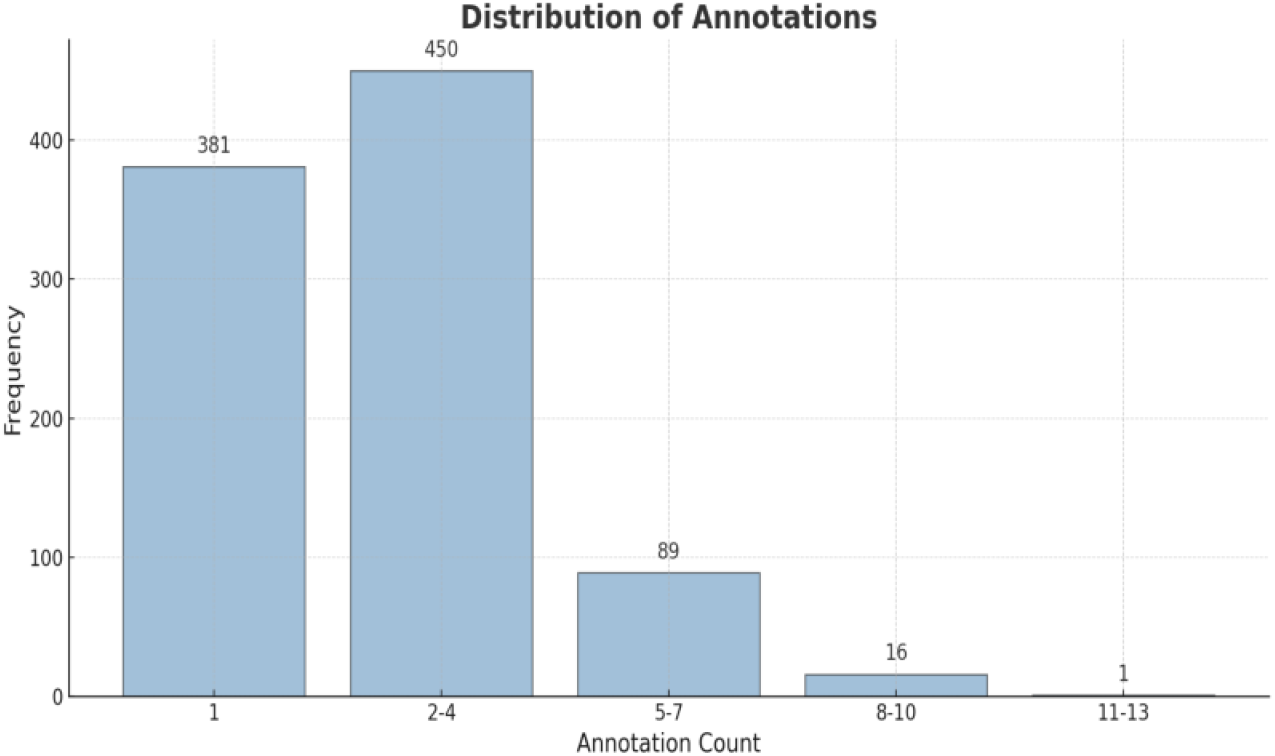
Distribution of the frequency of the number of annotated classes in the dataset.

### Dataset preprocessing and augmentation

A better recognition form the DL algorithm requires augmenting the information in the database. Images from the dataset, which are then divided into three subsets: training (70%), validation (10%), and test images (10%). Subsequently, images undergo a pre-processing phase, which includes auto-orientation and resizing to 640x640 pixels, with the objective of ensuring proper alignment and reducing computational load. Then data augmentation techniques were employed, including horizontal flipping and 90-degree rotations (clockwise, counter-clockwise, and upside down), with the objective of enhancing the diversity of the dataset and facilitating more effective model generalization. The augmented training set comprised 1,338 images, of which 130 were allocated for validation and 63 for testing.

### YOLOv8

The YOLO (You Only Look Once) model has made significant strides in the field of computer vision, representing a notable success. Developed by the Ultralytics company (13), it has gained widespread recognition. YOLOv8 boasts improved detection accuracy and speed, making it a leading choice for various applications in computer vision. This version of YOLO is employed for training purposes.

### Automated Counting Process

To enable rapid and precise scanning of the Sedgewick-Rafter counting chamber, we configured the software to define the coordinates for area acquisition and the reference image. This was achieved by identifying three reference points for each, corresponding to the minimum and maximum positions of the S1 and S2 motors along the X and Y axes. Based on these coordinates, the software calculates the number and positions of the scan fields needed to cover allthe specimen.

At the start position, the S1 and S2 servomotors automatically align with the upper-left corner of the area acquisition of the Sedgewick-Rafter chamber (indicated by the blue rectangle in Figure 6). The chamber is scanned in a snake-like pattern from left to right and top to bottom.

**Figure 6:**
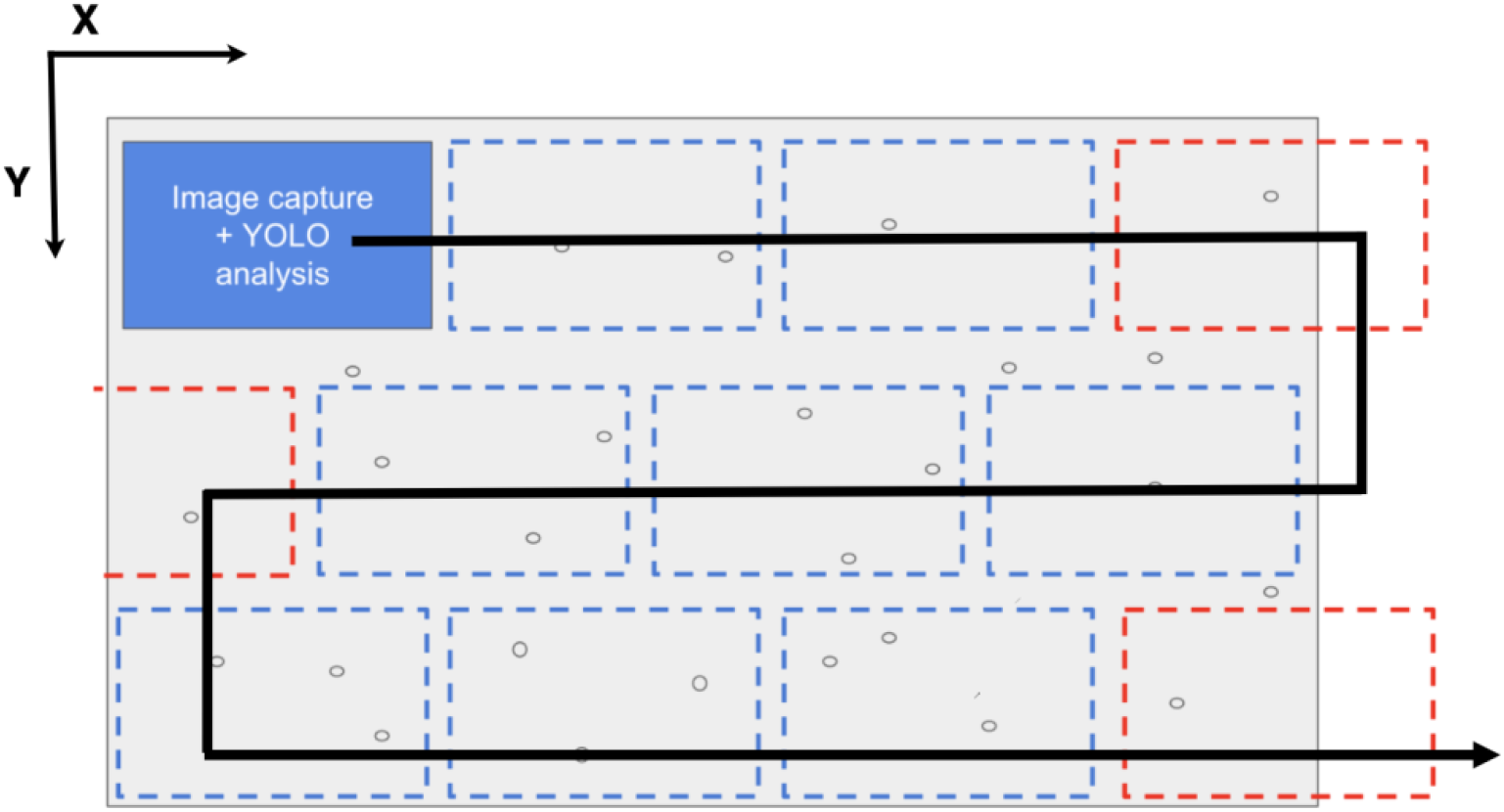
Camera trajectory to analyze the plate (black arrow). The dotted blue boxes represent the areas covered, i.e. viewed and captured by the camera. And the red ones represent the areas not covered entirely.

Next, a total of 39 images, each at a resolution of 1280×720 pixels, are captured at 1000× magnification (illustrated by blue dotted lines in Figure 6) to cover acquisition area. The number of rotifers in each image is recorded, and the population density is calculated by dividing the total number of rotifers by the number of images. This value is adjusted by the dilution factor of the rotifer sample and normalized using the image area volume of 45 μL. The volume is determined by calibrating the pixel dimensions of individual squares in the image against their physical size in the counting chamber (1 square (1mm^2^) = 1 μL).

## Results

The Rotiferometer’s performance was evaluated comprehensively, encompassing its accuracy, consistency, and utility in both experimental and practical applications. The subsequent section herein presents the outcomes emanating from four key evaluations: Model training results, Correlation between manual and automated counts using the Rotiferometer, Operator variability in rotifers counting using Rotiferometer. Then the Rotifers concentration dynamics over 40 days monitored using the Rotiferometer.

### Model training results

The training was performed in 100 epochs. The Table 3 presents the training results using various YOLOv8 models (YOLOv8n, YOLOv8s, YOLOv8m, YOLOv8l, and YOLOv8x) across several object detection metrics, as follows: the precision (P), recall (R), mAP@0.5, and mAP@0.5 : 0.95 values are presented. The values of losses, Precision, Recall, mAP and *F1-score* are obtained using the Equations respectively 1,2, 3 and 4.

**Table 3.**
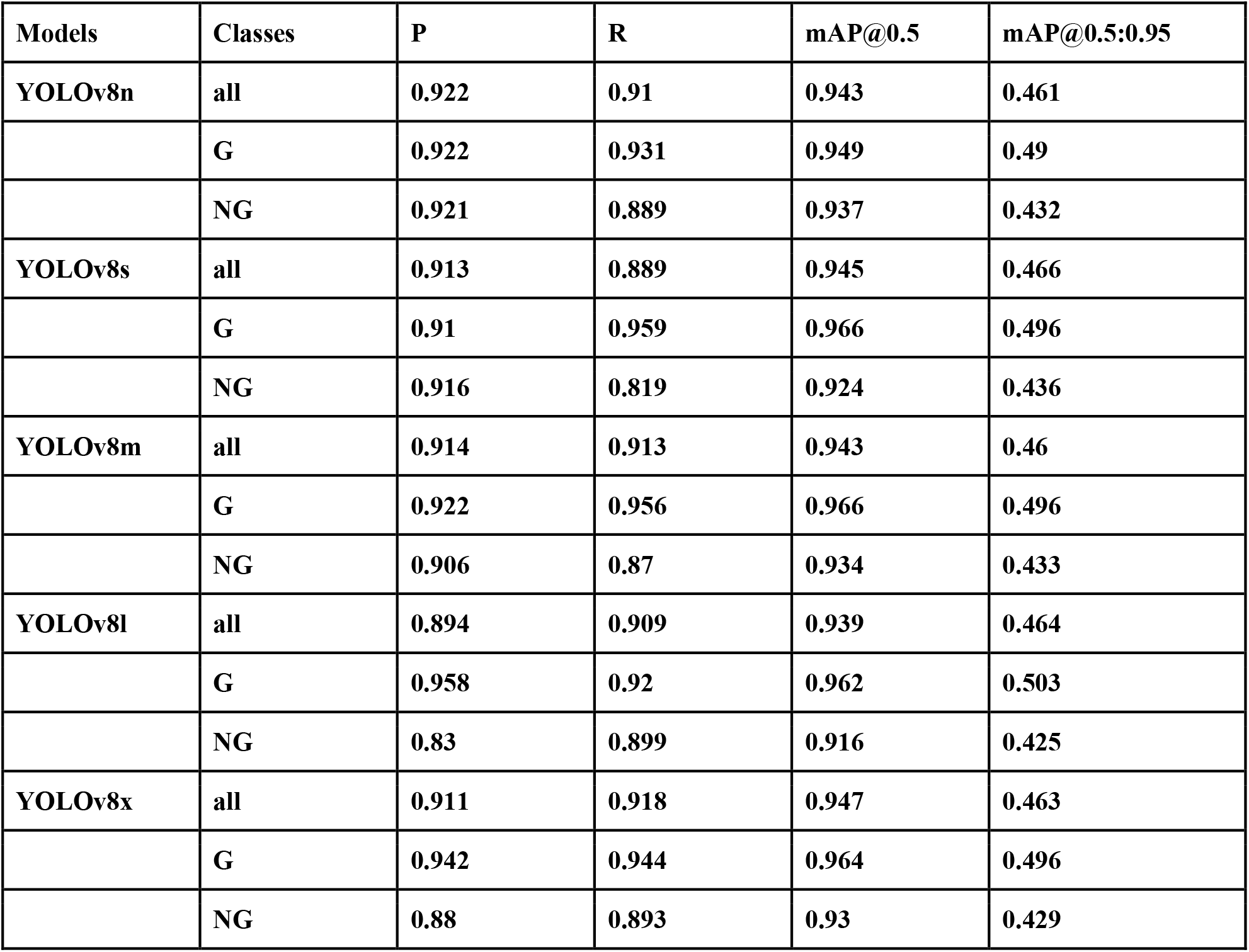
Training results for each model : Higher is better.

The *mAP@0*.*5* values are relatively high for all models, with a range of 0.911 to 0.947 (see Table 5), indicating that each model performs well for detecting objects at an IoU threshold of 0.5. In all trained models, the ′G′ class demonstrates greater stability across confidence levels, while the ′NG′ class exhibits a more pronounced decline, indicating that ′NG′ predictions are more challenging to classify with reliability and the ′G′ class possess more distinctive visual characteristics (carrying-egg), rendering them more readily discernible across models. The YOLOv8x model achieves the highest *mAP@0*.*5* score (0.947), closely followed by the YOLOv8s (0.945) and YOLOv8n (0.943) models. This indicates that the YOLOv8x model is the most precise, rendering it suitable for applications where accuracy is of paramount importance.

The *F1-score* (Figure 7.(a)) is paramount in object detection tasks, as it optimizes precision and recall. A high *F1-score* signifies that the model adeptly detects objects while minimizing false positives and false negatives. The graph illustrates that the model attains its maximum *F1-score* (0.91) at a confidence threshold of 0.483. Beyond this threshold, the score declines, suggesting that setting the confidence too high eliminates false positives but at the cost of missing actual detections (false negatives increase). This emphasizes the importance of threshold tuning in real-time detection of rotifers. The second graph of Figure 7 (Figure 7.(b)) displays the training and validation loss curves over 100 epochs. Loss functions measure the alignment of the model’s predictions with the ground truth, and their behavior over training provides insights into model generalization. Initially, both losses decrease rapidly, indicating effective learning. However, after a certain point, the validation loss stabilizes while the training loss continues to decrease, suggesting potential overfitting (i.e. the model memorizes the training data but struggles with unseen data). This highlights the need for early stopping, dropout, or other regularization techniques to maintain a balance between model complexity and generalization.

**Figure 7:**
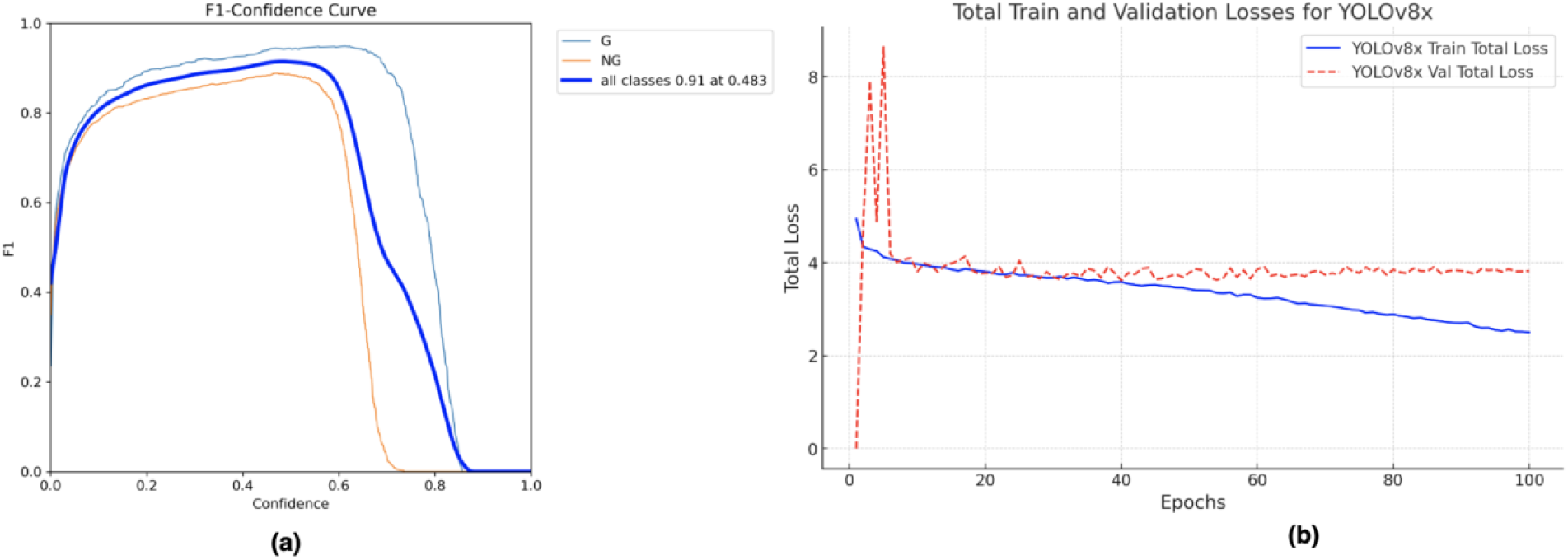
(a) : F1-score results for YOLOv8x. (b) : Validation loss and Training loss curves for YOLOv8x

When the same GPU was used for training, the model was tested on a dataset comprising 273 images, achieving an accuracy of 93.3% and a detection speed of 10.8 milliseconds per frame. As shown in Figure 8, the system is capable of detecting gravid and non-gravid rotifers using the trained model with YOLOv8x. This model (YOLOv8x) is then used for further experiments.

**Figure 8:**
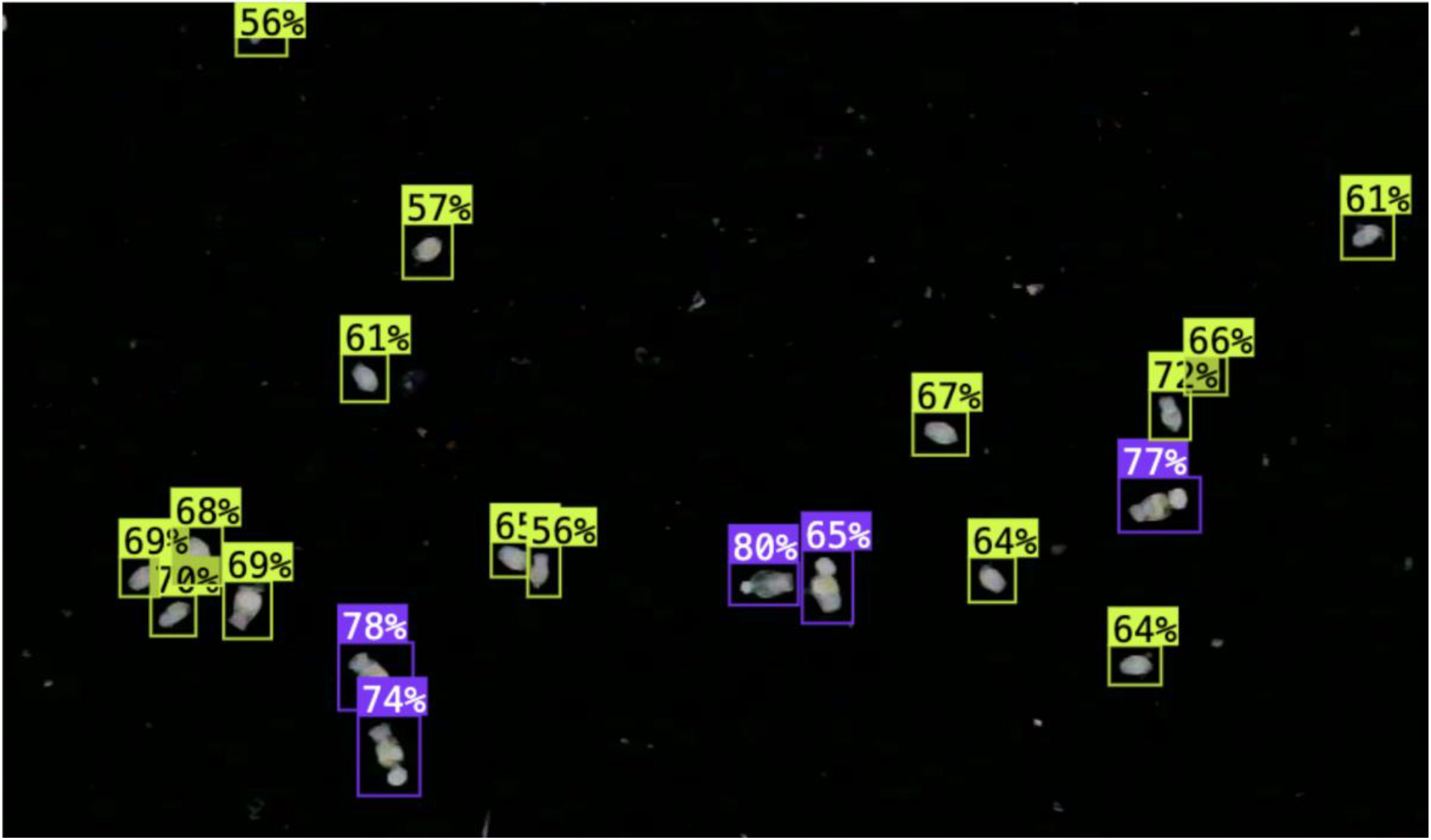
Example of detection using the trained model with YOLOv8x. Green color represents the ′*NG*′ class and the violet color is for the ′*G*′ class, with confidence of the model in percentage.

The Rotiferometer then was installed in the laboratory for technicians to conduct a series of validation tests over the course of several months.

### Evaluation of the accuracy of the Rotiferometer by comparison with manual counting method

We evaluated the accuracy of the Rotiferometer by comparing its results with those obtained by manual counting method for two categories of rotifers, gravid (with eggs) and non-gravid (without eggs). An experienced technician counted rotifers manually by observing the samples under a binocular microscope. Samples were prepared in 7 dilutions to simulate different rotifer concentrations (55-450 rotifers/ml). Each dilution was carefully prepared and the rotifers were manually counted to establish a reliable reference dataset.

As demonstrated in Figure 9.a, the comparison between manual and automatic counting for non-gravid rotifers is illustrated, with the regression equation being represented as follows: *Y* = 1.099*X* + 13.27. The regression coefficient, denoted by the slope of 1.099, indicates that the automated system overestimates the count by approximately 9.9 % compared to the manual method. Of greater significance is the intercept value of 13.27, which suggests the presence of a substantial systematic bias, indicating that even when the manual count approaches zero, the automated system still detects a certain number of non-gravid rotifers. This outcome signifies the existence of false positives, which are probably attributable to background noise, small debris, or the misclassification of overlapping structures as individual rotifers. The error bars are more pronounced in high-density samples, showing that as the concentration of non-gravid rotifers increases, the accuracy of the system declines. This finding suggests that in higher-density populations, the automated system may encounter difficulties in distinguishing individual rotifers, potentially resulting in object merging and over-counting. Despite the observed overestimation in high concentrations, the automated system appears to perform well at lower concentrations. When the manual count is small, the data points are closely aligned with the regression line, indicating a strong correlation between manual and automatic counting in these cases. The error remains minimal in this range, meaning that the system can accurately detect non-gravid rotifers in lower-density samples. This finding suggests that the algorithm effectively identifies and quantifies individual rotifers when they are more spatially separated, but as clustering increases, the accuracy decreases due to detection errors and segmentation difficulties. Figure 9.b, which represents the data for gravid rotifers, adheres to the regression equation: *Y* = 1.072*X* + 0.5917. slope of 1.072 indicates a minor overestimation of approximately 7.2 %, while the intercept value of 0.5917 is notably lower compared to the non-gravid scenario. This low intercept indicates minimal systematic bias, suggesting that the system does not introduce substantial errors when manual counts are low. The error bars remain small across all concentrations, showing that the system performs consistently well regardless of the number of gravid rotifers in the sample. The regression line is closely aligned with the data points, suggesting that the automatic system is highly reliable for detecting gravid rotifers, even in high-density samples.

**Figure 9.**
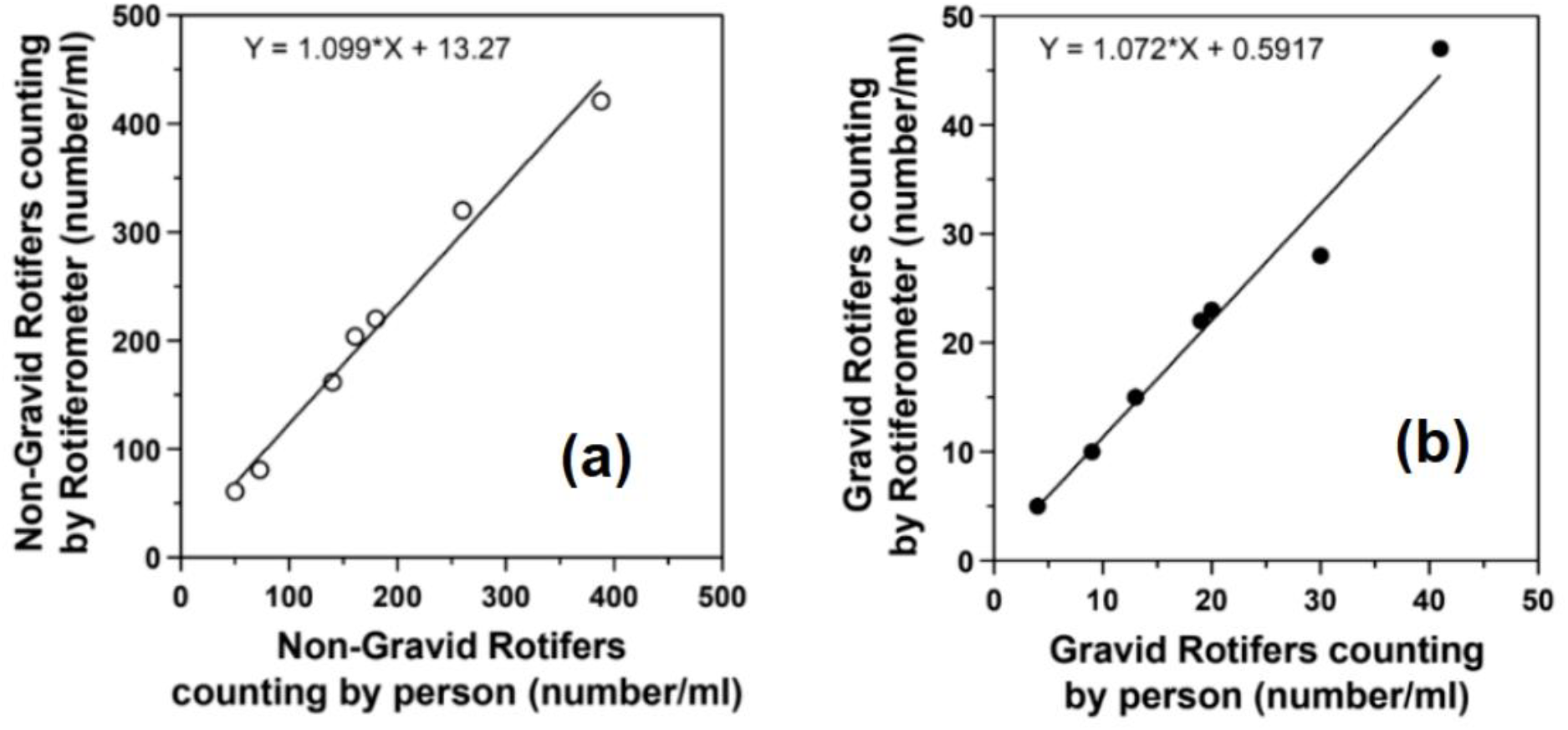
(a) : Linear regression of Rotiferometer vs. manual counts for Non-gravid Rotifers for 7 different cultures. (b) : Linear regression of Rotiferometer vs. manual counts for gravid Rotifers for 7 different cultures.

The comparison between the two graphs highlights a key distinction in the performance of the automated counting system depending on rotifer type and concentration. While the system demonstrates efficacy for both gravid and non-gravid rotifers at lower concentrations, it exhibits a tendency towards overestimation in higher-density samples, particularly for non-gravid rotifers. The significant intercept value in the non-gravid case suggests that the system has difficulty differentiating rotifers from background artifacts, leading to false positives. This effect is less pronounced for gravid rotifers, where the low intercept value confirms that false detections are minimal. The observed overestimation in high concentrations is likely due to increased clustering, making it harder for the system to accurately separate individual rotifers. This hypothesis is further substantiated by the error bars in the non-gravid dataset, which demonstrate increased variability as the number of rotifers rises. In contrast, for gravid rotifers, the detection accuracy remains stable, even at higher concentrations. This observation indicates that the algorithm is optimised for distinguishing gravid rotifers, potentially due to their more distinctive morphological characteristics. In contrast, non-gravid rotifers are more likely to be misclassified. The strong correlation (*R*^*2*^ values of 0.9729 and 0.9868 for gravid and non-gravid rotifers) observed in low-density samples indicates that when rotifers are well-separated, the system accurately detects and counts them. This suggests that the image processing algorithm performs optimally when objects do not overlap. However, as the density of rotifers increases, the Rotiferometer system’s performance deteriorates, resulting in elevated error rates and an overestimation of their numbers. It is evident that future refinements to the segmentation techniques may be beneficial in enhancing the system’s capacity to effectively detect and enumerate rotifers in high-density populations. Potential enhancements could involve the implementation of more sophisticated object separation methodologies or post-processing filters, which have the potential to mitigate false positives.

These findings confirm the hypothesis that the Rotiferometer system is a valuable tool for rotifer population quantification. However, it is clear that adjustments are needed to improve the system’s accuracy in high-density non-gravid samples. The strong agreement with manual counting in low concentrations suggests that the system is already effective in certain conditions. By refining the system’s detection thresholds and object differentiation capabilities, the overestimation in high-density samples could be mitigated, enhancing the automated method’s precision and reliability.

### Evaluation of the Reproducibility and Repeatability of Rotifer Counts Using the Rotiferometer

We evaluated the reproducibility and repeatability of the Rotiferometer. Two operators, designated A and B, independently analyzed the same rotifer sample three times (1-3), as shown in Figure 10. To assess the reproducibility, we compared the recorded counts between the two operators. The mean with standard deviation for the gravid rotifers (from operator A: 2.62 × 105 ± 1.34 × 104 and operator B: 2.60 × 105 ± 1.88 × 104) and non-gravid rotifers (from operator A: 8.50 × 105 ± 1.11 × 104 and operator B: 8.27 × 105 ± 2.44 × 104) showed no significant differences between the operators’ results. The percentage coefficient of variation (% CV) was calculated for both operators, with values ranging from 5–7% for non-gravid rotifers and 1–3% for gravid rotifers. These results fall within the acceptable range of 0–10%, indicating a high level of repeatability for the Rotiferometer. These results confirm that the Rotiferometer provides consistent and reproducible measurements for both gravid and non-gravid rotifers, regardless of the operator performing the analysis.

**Figure 10.**
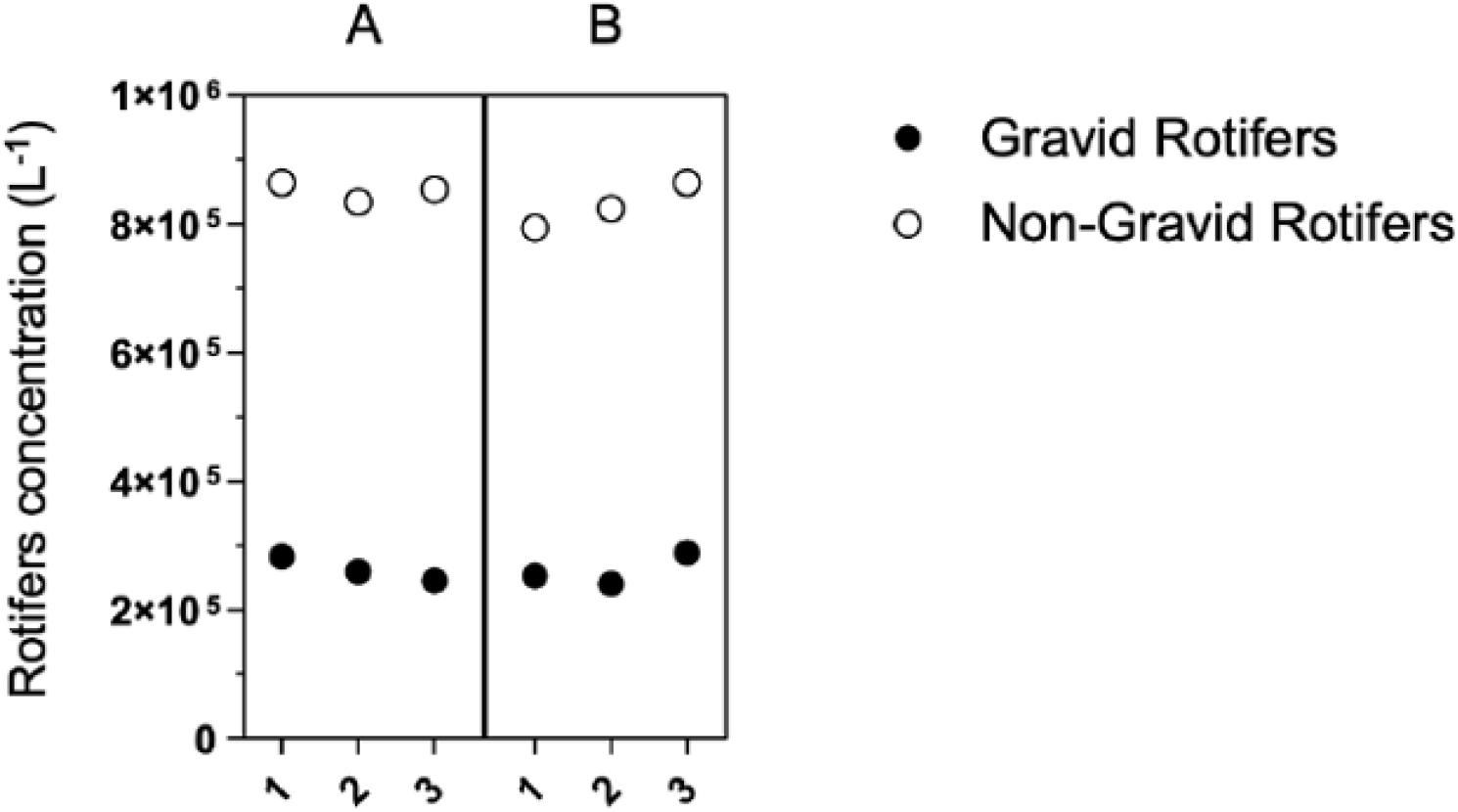
Variability in rotifer counting between operators A and B across the triplicates (1-3), using the Rotiferometer. non-gravid rotifer counts are represented by filled circles, while gravid rotifer counts are depicted by empty circles.

### Real-Time Monitoring of Rotifer Culture Dynamics

We investigated whether the Rotiferometer could provide a consistent approach to monitoring rotifer culture. We tested the device over 45 days under conditions that replicated standard rotifer culture practices used in aquaculture. Daily measurements of both gravid and non-gravid rotifer concentrations were obtained using the Rotiferometer. The gravid rotifer is a well-known feature of growing rotifers. The eggs of these rotifers will hatch and replenish the culture and can therefore be used as an indicator of culture growth. The total number of rotifers was plotted over time with the percentage of gravid rotifers superimposed on the growth curve to visualise the population dynamics (Figure 11).

**Figure 11.**
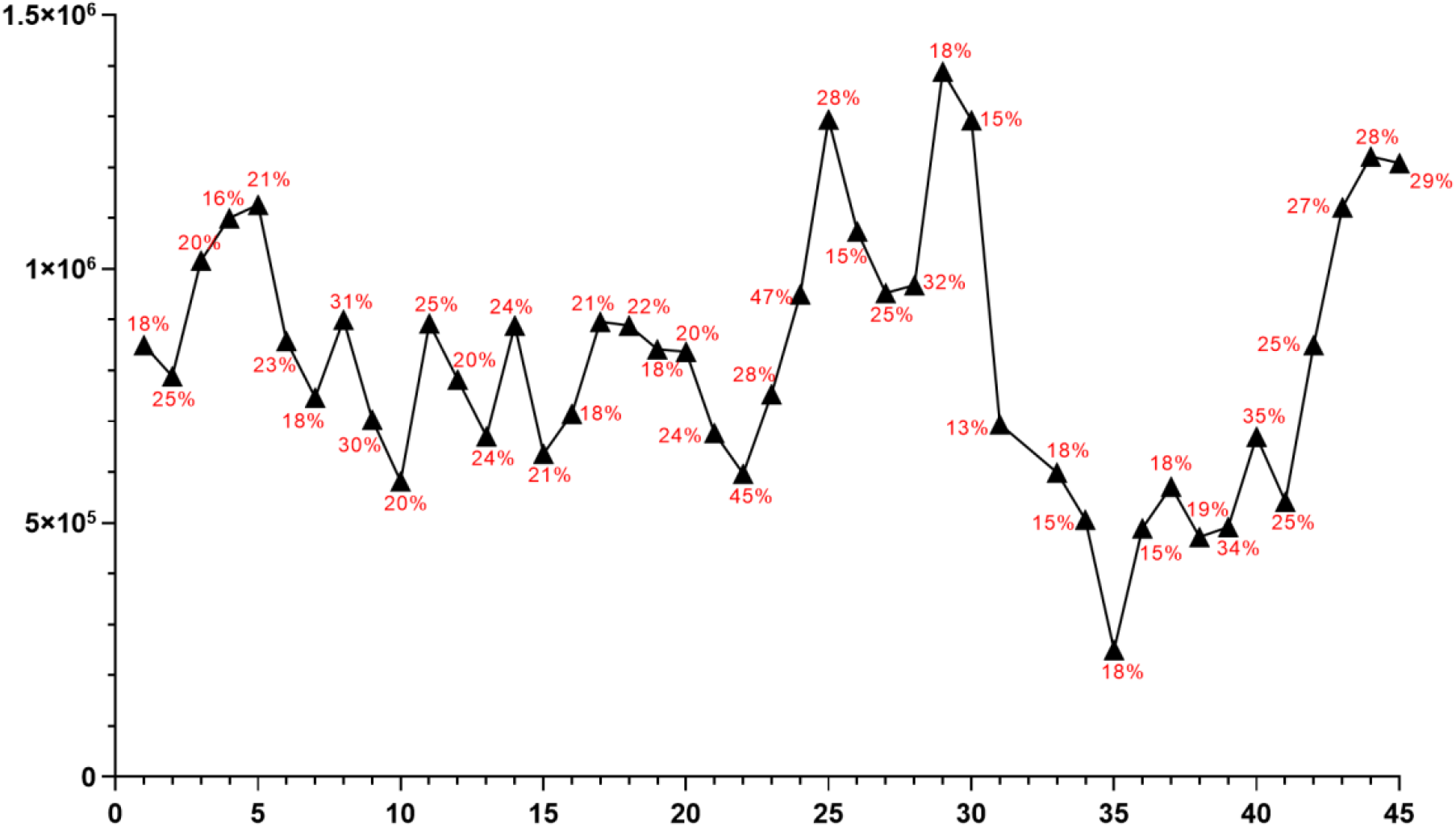
Rotifers concentration dynamic over 45 days monitored using the Rotiferometer.

We started the rotifer culture with an expansion phase (day 0-5), which aimed to increase the number of rotifers by successive additions of culture medium. The culture was started with a rotifer density of 8 × 105 per litre and 18% gravid rotifers in a 4 litre culture (Figure 11). Over the next five days, the volume was gradually increased by 1 litre per day, reaching 8 litres on day 5. As expected, analysis of the resulting culture showed an increased number of rotifers during the expansion phase, reaching a peak population of 1.1 × 106 total rotifers with 21% gravid rotifers on day 5 (Figure 11). Following the expansion phase, we initiated the harvest and maintenance phase (days 5-35). During this period, 20-30% of the rotifers were harvested daily to feed the fish larvae, while 30% of the culture medium was replaced to maintain a constant volume of 8 litres. As shown in Figure 11, we observed two main trends in the rotifers culture during this period. A first trend occurred between days 5 and 29, when the rotifer density oscillated between minimum and maximum values of 5.8 10^5^-1.3 10^6^ rotifers and 15-47% gravid rotifers, reflecting a dynamic equilibrium between growth and harvesting of rotifers. The second trend occurred when the density of rotifers gradually decreased from day 29 (1.3 10^6^ rotifers) to day 35 (2.5 10^5^ rotifers). This decrease was attributed to the aeration problem in the rotifer culture, which led to oxygen depletion and subsequent rotifer mortality. To increase rotifer numbers, we reinitiated the expansion phase between days 35 and 39 by suspending rotifer harvesting while maintaining 30% daily media renewal and gradually increasing the culture tank volume to a final capacity of 8 litres. Despite these efforts, the rotifer density remained low and did not exceed 5.7 10^5^ rotifers. Consequently, from day 39 to day 45, we continued the 30% daily media renewal without further increasing the culture volume or harvesting the rotifers. This approach allowed the rotifer population to increase and stabilise at a density of 1.2 × 106 rotifers per litre, with 29% gravid individuals.

These results highlight the effectiveness of the Rotiferometer in facilitating real-time observation of rotifer culture dynamics, allowing identification of critical phases and timely intervention to ensure optimal culture conditions.

## DISCUSSION

These results validate the system’s ability to perform reliable real-time quantification of rotifers while eliminating operator counting process. However, discrepancies observed in the vicinity of image edges or with overlapping individuals underscore the necessity for further refinement of the image processing algorithms. Notwithstanding, the system’s capacity to store and export data in real-time, coupled with automated generation of population dynamics graphs, provides a significant advantage over traditional manual methods. Furthermore, the Rotiferometer system is capable of scanning a 1 *ml* plate within 3 minutes, thereby providing results. Conversely, manual enumeration of rotifers typically requires a time frame of 3 - 20 minutes, contingent upon the density of rotifers within the sample. Consequently, the Rotiferometer is six times faster than manual counting for high-density samples. The system’s robustness is further confirmed in the second experiment, which evaluated operator variability. Measurements performed by different operators showed no significant differences, with consistent mean and standard deviation values for both gravid and non-gravid rotifers. This consistency underscores the reliability of the Rotiferometer and reduces the need for extensive operator training, making it a valuable tool for environments where multiple personnel handle rotifer culture monitoring. The integration of advanced DL models, specifically YOLOv8, into the system serves to further enhance its functionality. YOLOv8 achieved high precision and recall rates, with the best model reaching a mAP of 94.7% for classifying rotifers. This capability enables the Rotiferometer to accurately differentiate between gravid and non-gravid rotifers, a critical factor for assessing the health and productivity of cultures. Furthermore, the system’s capacity to automate repetitive tasks, including counting, data recording and graph generation, has been demonstrated to significantly reduce labor demands, thereby enabling technicians to focus on more strategic management tasks. Despite its many advantages, certain limitations were identified. For example, edge effects and occasional misclassification of overlapping rotifers remain challenges. Future work should focus on improving the image processing pipeline, enhancing the YOLOv8 model with additional training data, and incorporating features to address edge cases. A critical aspect of enhancing object detection accuracy is the process of hyperparameter tuning. A further disadvantage is that the operator must eliminate the rotifers with vinegar in order to quantify them. While it is acknowledged that the number of rotifers eliminated is negligible in comparison to the culture, it is nevertheless preferable to avoid this step.

## CONCLUSION

The Rotiferometer has been developed as a robust, integrated solution for the automated monitoring and quantitative analysis of rotifer cultures. This addresses a critical need in both aquaculture production systems and experimental research environments. The platform is developed to address the limitations of manual rotifer monitoring techniques by unifying cost-efficient mechatronic engineering, high-contrast transmitted light microscopy, and deep learning-based image analysis. A salient feature of the Rotiferometer is its operational simplicity, which facilitates its deployment by non-specialized personnel in routine monitoring workflows. This renders the system especially well-suited for aquatic animal facilities, live feed hatcheries, and laboratories engaged in rotifer-related experimental studies. As demonstrated in the Supplementary material (video S1), the system is straightforward to use and requires minimal setup, which supports its potential for wide adoption in operational settings. The system has been developed for the precise and high-throughput quantification of rotifers. It employs an automated, calibrated two-axis motion platform that enables the scanning of a standard Sedgewick-Rafter chamber. It utilise a deep learning detection pipeline based on YOLOv8 to perform two functions. Firstly, it counts individual rotifers. Secondly, it distinguishes between gravid and non-gravid individuals. The latter is a critical metric for evaluating the reproductive status and productivity of the culture. It is imperative to recognise the significance of this functionality in the context of longitudinal assessment of cultural health, as it facilitates the timely identification of any fluctuations in reproductive output. In comparison with conventional methods, the automated counting process has been shown to significantly reduce human error and inter-operator variability, while concomitantly achieving a substantial enhancement in counting speed, reproducibility, and temporal resolution. Each scan session is meticulously documented, resulting in the generation of archived image datasets and quantitative outputs with precise date and time stamps. Digital traceability facilitates the monitoring of culture dynamics over time, thereby supporting both operational management and retrospective analysis. Another notable advantage of the system is its hardware and software modularity, which allows for future extension to other zooplankton species exceeding 100 μm in size. It is evident that the Rotiferometer architecture is characterised by a high degree of adaptability, which can be attributed to its capacity to adjust optical magnification, detection parameters and training datasets. This adaptability renders the Rotiferometer suitable for a range of applications, including copepod monitoring, larval quality assessment and toxicological bioassays involving planktonic organisms. In summary, the Rotiferometer provides a scalable, user-friendly, and scientifically rigorous platform for automated rotifer monitoring. The system combines the qualities of speed, accuracy, and reproductive-state discrimination with archiving capabilities and extensibility, positioning it as a key enabler of data-driven management in both aquaculture and rotifer-based biological research. Its integration into existing workflows has the potential to significantly enhance live feed productivity, optimize animal growth conditions, and standardize rotifer culture protocols across laboratories and production facilities.

## Author contributions

All authors have made substantial contributions to the conception and design of the study. All authors contributed on images acquisition and analysis, labeling images (AM, MA), development of the deep learning model (AD), automation process workflow (DN, LB), mechanical design (LB, SH) and (AD, AM) wrote the main manuscript text and final approval of the version submitted (all authors).

## Acknowledgments

The authors thank Edouard Manzoni and Lois Bernabé technicians for the Animal Facility and Engineering Aquatic Models Platform of Sorbonne University for their help during experimental tasks.

